# Novel insights into phage biology of the pathogen *Clostridioides difficile* based on the active virome

**DOI:** 10.1101/2023.09.27.559748

**Authors:** Miriam A. Schüler, Rolf Daniel, Anja Poehlein

## Abstract

The global pathogen *Clostridioides difficile* is a well-studied organism, and researchers work on unravelling its fundamental virulence mechanisms and biology. Prophages have been demonstrated to influence *C. difficile* toxin expression and contribute to the distribution of advantageous genes. All this underlines the importance of prophages in *C. difficile* virulence. Although several *C. difficile* prophages were sequenced and characterized, investigations on the entire active virome of a strain are still missing. Phages were mainly isolated after mitomycin C-induction, which does not resemble natural stressor for *C. difficile.* We examined active prophages from different *C. difficile* strains after cultivation in the absence of mitomycin C by sequencing and characterization of particle-protected DNA. Phage particles were collected after standard cultivation, or after cultivation in the presence of the secondary bile salt deoxycholate (DCA). DCA is a natural stressor for *C. difficile* and a potential prophage-inducing agent. We also investigated differences in prophage activity between clinical and non-clinical *C. difficile* strains. Our experiments demonstrated that spontaneous prophage release is common in *C. difficile*, and that DCA presence induces prophages. Fourteen different, active phages were identified by this experimental procedure. We could not identify a definitive connection between clinical background and phage activity. However, one phage exhibited distinctively higher activity upon DCA-induction in the clinical strain than in the corresponding non-clinical strain, although the phage is identical in both strains. We recorded that enveloped DNA mapped to genome regions with characteristics of mobile genetic elements other than prophages. This pointed to mechanisms of DNA mobility that are not well-studied in *C. difficile* so far. We also detected phage-mediated lateral transduction of bacterial DNA, which is the first described case in *C. difficile*. This study significantly contributes to our knowledge on prophage activity in *C. difficile* and revealed novel aspects on *C. difficile* (phage) biology.

## Introduction

The pathogen *Clostridioides difficile* significantly contributes to nosocomial infections worldwide (1). A *C. difficile* infection mainly establishes after antibiotic treatment due to diverse resistances in *C. difficile* strains combined with the disturbed intestinal microflora (2). An intact microbiome usually provides resistance to *C. difficile* colonization and disease manifestation by producing different secondary bile salts such as lithocholate and deoxycholate (3). These compounds impede *C. difficile* spore germination and cell growth (4). Symptoms of an established *C. difficile* infection are caused by its toxins, predominantly toxin A and B, which are encoded in the pathogenicity locus (5). Symptom severity ranges from mild to severe manifestation, which also could include death of the infected individual (1). The personal health condition significantly affects resistance against a *C. difficile* infection, but the infecting strain is of relevance as well (6). Different *C. difficile* strains are linked to divergent virulence, and various aspects such as toxin production levels or secondary bile salt resistance were shown to correlate with disease severity (7,8). However, these studies also partially contradict in the concluded relevance of specific features on virulence. In addition to general virulence factors, mobile genetic elements (MGE) also contribute to *C. difficile* virulence (9–12). Those elements play a key role in fast adaptation to environmental conditions via horizontal gene transfer (HGT) (13). *C. difficile* genomes harbor various MGEs, including prophages (14). Prophages are widespread among the species *C. difficile*, and multiple prophages can exist within the same host (15). Primarily, prophages were assumed to affect *C. difficile* virulence solely by encoding advantageous traits such as antibiotic resistances, which was demonstrated by phage-mediated transduction of an erythromycin resistance (10).Phages can influence toxin production in *C. difficile* strains (11,12). These findings drew further attention to the influence of phages on *C. difficile* virulence. Phage research commonly works with prophage induction by introducing stressors such as UV radiation or mitomycin C. Meanwhile, studies confirmed spontaneous prophage release from *C. difficile* isolates, and clinically relevant antibiotics were also investigated for their phage-inducing effect (16,17). All experiments on *C. difficile* phages however worked with cultivation conditions that do not represent the actual habitat. Some components of the intestinal environment are stressful for *C. difficile*, such as the secondary bile salts. One of the prominent secondary bile salts is deoxycholate (DCA), which can promote biofilm formation in *C. difficile* (18). It was further demonstrated that DCA induces the bacterial SOS response (19). Activation of the SOS response in turn induces prophages and lead to phage particle production and release via host cell lysis (20). It is therefore likely that DCA induces prophages as well, which would be a critical aspect in *C. difficile* biology and shed new light on genetic transfer *in vivo*.

In this work, we examined prophage activity in different *C. difficile* strains under spontaneous conditions and in the presence of DCA. The analyzed *C. difficile* strains were of non-clinical and clinical origin and corresponded pairwise to each other based on their sequence type (ST). Active prophage regions in these strains were identified by sequencing of particle-protected DNA, analyzed for DCA-induced activity and for possible differences between clinical and non-clinical strains that could contribute to virulence. The sequencing approach is more sensitive than electron microscopy or plaque assays, the commonly used detection methods for active *C. difficile* phages. We could therefore detect active prophages that might otherwise be missed due to insufficient activity, but could contribute to HGT.

## Methods

### Strains and cultivation conditions

The *C. difficile* strains used in this study were of clinical or non-clinical clinical background (personal communication), with four pairs of clinical and non-clinical strains correpsonding in ST, and one additional non-clinical strain (Table 1). Strains were routinely cultivated under anaerobic conditions in supplemented Brain Heart Infusion Broth (BHIS; supplemented with 0.5% yeast extract, 0.05% L-cystein, 0.0001% Na-resazurin, purged with nitrogen) at 37°C. Putative prophage regions of the strains were predicted with PHASTEST (21).

**Table 1.** *C. difficile* strains used in this study. Strain information on ST (Clade), profile of toxins A. B, and CDT, clinical or non-clinical background, and GenBank accession of the genome is listed.

| Strain | ST (Clade) | Toxin profile | Background | GenBank accession |
| --- | --- | --- | --- | --- |
| TS3_3 | 1 (2) | A<sup>+</sup>B<sup>+</sup>CDT<sup>+</sup> | Non-clinical | CP134872 |
| DSM 28196 | 1 (2) | A<sup>+</sup>B<sup>+</sup>CDT<sup>+</sup> | Clinical | CP012320.1 |
| B1_2 | 3 (1) | A<sup>+</sup>B<sup>+</sup>CDT<sup>-</sup> | Non-clinical | CP132141-43 |
| SC084-01-01 | 3 (1) | A<sup>+</sup>B<sup>+</sup>CDT<sup>-</sup> | Clinical | CP132146-48 |
| J2_1 | 8 (1) | A<sup>+</sup>B<sup>+</sup>CDT<sup>-</sup> | Non-clinical | CP134690-1 |
| SC083-01-01 | 8 (1) | A<sup>+</sup>B<sup>+</sup>CDT<sup>-</sup> | Clinical | CP132144-45 |
| MA_1 | 11 (5) | A<sup>+</sup>B<sup>+</sup>CDT<sup>+</sup> | Non-clinical | CP132139-40 |
| DSM 29747 | 11 (5) | A<sup>+</sup>B<sup>+</sup>CDT<sup>+</sup> | Clinical | CP019864.1 |
| MA_2 | 340 (C-III) | A<sup>-</sup>B<sup>-</sup>CDT<sup>-</sup> | Non-clinical | CP129431-32 |

### DCA tolerance assessment

The stress effect of DCA on the various strains was assessed via a minimum inhibitory concentration (MIC) assay and relative growth determinations at different concentrations. Cultivation was performed in cell culture plates (24 well for suspension cells, Sartorius AG, Göttingen, Germany) in an anaerobic tent (Coy Laboratory, Grass Lake, USA). Overnight cultures of *C. difficile* strains were cultivated as described above and used to inoculate two ml medium to a final OD_600_ of 0.05, with either BHIS medium alone or supplemented with DCA (sodium deoxycholate, Sigma-Aldrich Chemie Gmbh, Taufkirchen, Germany) in concentrations ranging from 0.255 mM to 1.2 mM, which cover the physiological concentration range in humans (22). Culture plates were incubated at 37°C for 22 hrs and kept anaerobic during the OD_600_ measurement in a Synergy 2 Plate Reader (Biotek Agilent Technologies Deutschland GmbH, Böblingen, Germany). Relative growth in the presence of DCA was calculated in relation to the untreated cultures. Each experiment was performed in triplicates. Relative growth was visualized with ggplot2 (v3.4.2) (23) in RStudio (v2022.06.0) (24) and significance determined with the Tukey’s ‘Honest Significant Difference’ method implemented in the stats package (v4.2.0) (25).

### Prophage induction and phage particle isolation

Phage particles were isolated from untreated and DCA-induced cultures. Two pre-warmed flasks of 55 ml BHIS for each isolate were inoculated 1:100 from an overnight culture and incubated at 37°C. Growth was monitored via OD_600_ measurements until an OD of ∼0.6 (0.5-0.7). One culture for each isolate was induced with 0.255 mM (0.01%) DCA final concentration. The physiological concentration of DCA varies between individuals (22). We therefore used this concentration that is within the physiological range and was also used in various *C. difficile* studies regarding growth behavior or spore germination (7,26). The DCA solution was freshly prepared under anaerobic conditions, with DCA suspended in distilled water so that 100 µl inducing solution was required per 10 ml culture. The solution was sterilized by filtration (Filtropur S 0.2 µm, Sarstedt AG & Co. KG, Nürnbrecht, Germany) and anaerobically added to the induction cultures. The second culture was not induced for analysis of spontaneous phage activity. Induced and non-induced cultures were further incubated until 22 hrs total incubation. The final OD_600_ of each culture was determined before isolating phage particles.

For phage particle isolation, cells were pelleted via centrifugation at 4°C and 3,000 x *g* for 15 minutes. Remaining cells were removed by filtration of the supernatant with a 0.45 µm Filtropur S filter (Sarstedt). Phage particles were pelleted via centrifugation at 8°C and 20,000 x *g* for 1 h. The pellet was suspended in 1 ml SM buffer (50 mM Tris-HCl, 100 mM NaCl, 8 mM MgSO_4_ · 7 H_2_O, pH 7.4) supplemented with 0.5 mM CaCl_2_ (supporting phage stability and upcoming DNase treatment) and let soak overnight at 4°C. Particle suspension was further supported the next day by shaking at 150 rpm (LT-V Lab-Shaker, Adolf Kühner AG, Birsfelden, Germany) at room temperature for 2 hrs. Suspended samples were finally transferred to 2 ml DNA LowBind micro tubes (Sarstedt) for following treatments using cut filter tips to reduce possible shearing. Samples were stored at 4°C.

### Isolation of particle-protected DNA from phage samples

Phage DNA was isolated using the MasterPure Gram Positive DNA Purification Kit (Epicentre, Madison, WI, USA) with modifications. Prior to the phage DNA isolation, phage samples were supplemented with 2 µl of 100 mg/ml lysozyme solution (lysozyme from chicken egg white 177,000 U/mg; Serva, Heidelberg, Germany) suspended in SM buffer to remove remaining host cell debris, and with 50 µg/ml final concentration RNase A (Biozym Scientific GmbH, Hess. Oldendorf, Germany) and 10 U Baseline-ZERO DNase (Biozym Scientific GmbH) to digest host nucleic acids. Samples were incubated at 37°C for 6 hrs with gentle inversion every 30 minutes. Fragments of host nucleic acids resulting from the digestion were removed by phage particle pelleting via centrifugation at 4°C and 20,000 x *g* for 1 hour. The recovered pellet was suspended in 150 µl SM buffer. Suspension was supported by shaking at 150 rpm (LT-V Lab-Shaker) at room temperature for 10 minutes and slight flicking. EDTA (0.5 M, pH 8.0) was added to 10 mM final concentration for complete DNase inhibition.

Phage particles were lysed by adding 1% SDS (10% solution) and 2 µl Proteinase K (50 µg/µl; Biozym Scientific GmbH), incubation at 56°C for 1.5 hrs and gentle inversion every 30 minutes. Subsequently, samples were completely cooled down on ice before addition of 130 µl MPC Protein Precipitation Reagent (pre-cooled to −20°C). After mixing by gentle inversion, proteins were pelleted via centrifugation at 4°C and 10,000 x *g* for 10 minutes. The DNA-containing supernatant was transferred to 1.5 ml DNA LowBind micro tubes (Sarstedt). DNA was precipitated by addition of 0.3 M Na-acetate (3 M, pH 5.2), 10 mM MgCl_2_ (2 M), and 0.8 volume isopropanol at room temperature. Samples were inverted 40 times and incubated at room temperature for 10 minutes before centrifugation at 4°C and 15,000 x *g* for 30 minutes. The supernatant was removed carefully and the DNA pellet washed twice with 150 µl 70% ethanol (pre-cooled to −20°C) and centrifugation at 4°C and 15,000 x *g* for 5 to 10 minutes. The supernatant was removed and the sample briefly centrifuged again to collect all residual ethanol for final removal. DNA pellets were air-dried under a sterile bench and immediately suspended in 20 µl TE buffer. Complete DNA elution was supported by brief storage at 4°C and slight flicking, before final storage at −20°C. DNA concentration was assessed with the Qubit 3.0 Fluorometer (Thermo Fisher Scientific, Waltham, MA, USA) using the HS dsDNA assay kit.

### Phage DNA sequencing and sequencing read processing

Phage DNA was subjected to Illumina sequencing for dsDNA by paired-end library preparation with the Nextera XT DNA Library Preparation Kit (Illumina, San Diego, CA, USA) as recommended by the manufacturer. Libraries were sequenced using an Illumina MiSeq system and MiSeq Reagent Kit version 3 (2 × 300 bp, 600 cycles) as recommended by the manufacturer.

All following software was used in default mode. Sequencing raw reads were quality processed with fastp (v0.23.4) (27) before removing the sequencing adapters with Trimmomatic (v0.39) (28). Processed reads were mapped onto the corresponding host genome using bowtie2 (v2.5.0), and the resulting SAM file was converted to a TDS file for bioinformatics analysis with the TraV software (29).

### Data analysis of phage sequencing reads

TDS files of processed reads for the same isolate were together inspected in TraV (29) for read coverage, and reads were normalized by calculation of nucleotide activity per kilobase of exon model per million mapped reads (NPKM). This results in a value for each CDS corresponding to its read coverage in reference to the overall read amount. NPKM values were further normalized to account for the differing growth behavior under the induction conditions by transforming values to an OD_600_ of 2.0. In this way, NPKM values reflected phage abundance under the different conditions at same cell density, which allows a better qualitative estimation of phage activity. OD normalization and visualization of NPKMs values were done in Rstudio (v2022.06.0) (24) using the packages tidyverse (v2.0.0) (30), ggforce (v0.4.1) (31), and ggplot2 (v3.4.2) (23). NPKM values were plotted against the host genome with regard to sequence start of the corresponding CDS, and original and normalized NPKM values were plotted together to visualize the effect of OD normalization. Phage regions predicted with PHASTEST (21) were also implemented.

### Phage genome annotation and gene content analysis

Active regions identified via sequence read mapping were extracted for anew genome annotation. Sequence ranges were thereby adopted from PHASTEST (21) predictions or selected based on read mapping in TraV (29), if the predicted prophage region did not cover the entire mapped region. Annotation was customized for phage genomes using Pharokka (v1.3.2) (32) in default mode with sequence re-orientation to the large terminase subunit. If the large terminase subunit was not annotated automatically, it was determined via BLAST analysis (33) and manually annotated. For specific genes and their encoded protein, additional analyses with protein BLAST (33) and InterProScan (v5.63-95.0) (34) were performed.

### Phage genome-based classifications

Genome-based classifications were done with different bioinformatic analysis tools. An average nucleotide identity analysis (ANI) with pyani (v0.2.12) (35) and MUMmer3 alignment (36) (ANIm) was used in default mode to compare the phages amongst each other. Assessment of the DNA-packaging strategy was performed based on the study of Rashid et al. (37). The large terminase subunit was aligned at protein sequence level with ClustalW and a Maximum-Likelihood phylogenetic tree was constructed with the Whelan and Goldman (WAG) substitution model and otherwise default parameter with the software MEGA (v11.0.13) (38). Branches were collapsed in MEGA (38) if none of our phages clustered within and final modifications for visualization were done in Inkscape (v0.48; https://inkscape.org/de/). A nucleotide BLAST analysis (33) with default parameters was performed to check for similarity to already known phage genomes and for the prevalence in genomes of other *C. difficile* strains. BLAST results were ordered based on query coverage and hits with a query coverage below 10% were neglected, unless relevant matches with higher coverages were not obtained. For assigning the phages to a morphological family of the order *Caudovirales*, phage genomes were inspected for the presence of baseplate proteins and sequence length of the tail length tape measure protein (39). If the tail length tape measure protein was not annotated, it was identified via protein BLAST analysis (33) and manually curated in the genome.

### Data availability

The sequencing raw data is deposited at the NIH short read archive (SRA) under accessions listed in Table 2.

**Table 2.** SRA accessions of sequencing raw data.

| Strain | Spontaneous | DCA-induced |
| --- | --- | --- |
| TS3_3 | SRR26060497 | SRR26060498 |
| DSM 28196 | SRR26060430 | SRR26060431 |
| B1_2 | SRR26060462 | SRR26060463 |
| SC084-01-01 | SRR26064407 | SRR26064408 |
| J2_1 | SRR26060485 | SRR26060486 |
| SC083-01-01 | SRR26060487 | SRR26060488 |
| MA_1 | SRR26060489 | SRR26060490 |
| MA_2 | SRR26064320 | SRR26064321 |

## Results and discussion

### DCA-tolerance is linked to the genetic but not clinical strain background

Before investigating the effect of the secondary bile salt DCA on prophage activity in *C. difficile*, individual DCA tolerance of the various strains was assessed in form of a MIC assay with relative growth determinations (Fig 1). DCA concentrations ranged from 0.255 mM to 1.2 mM, thereby comprising the physiological human concentration of DCA (22). All strains already exhibited reduced growth at the lowest concentration, which further decreased with increasing concentration. At all concentrations, strains of the same ST showed no significant difference in DCA tolerance, which implied comparable stress levels. Similar stress levels in turn might imply similar DCA-induced prophage activity. In contrast, ST-specific tolerance differences were apparent across the DCA concentration range, with ST1 exhibiting highest tolerance, followed by ST8, ST11, ST3, and lastly ST340, which was the most susceptible ST. Consequently, DCA tolerance correlated with the ST but not with the clinical background. Tolerance difference between the STs was most distinct at the lowest concentration, which was also used in the prophage induction experiments. Determined MICs ranged from 1 mM (ST11 and ST340) to 1.2 mM (ST1, ST3, ST8).

**Fig 1.**
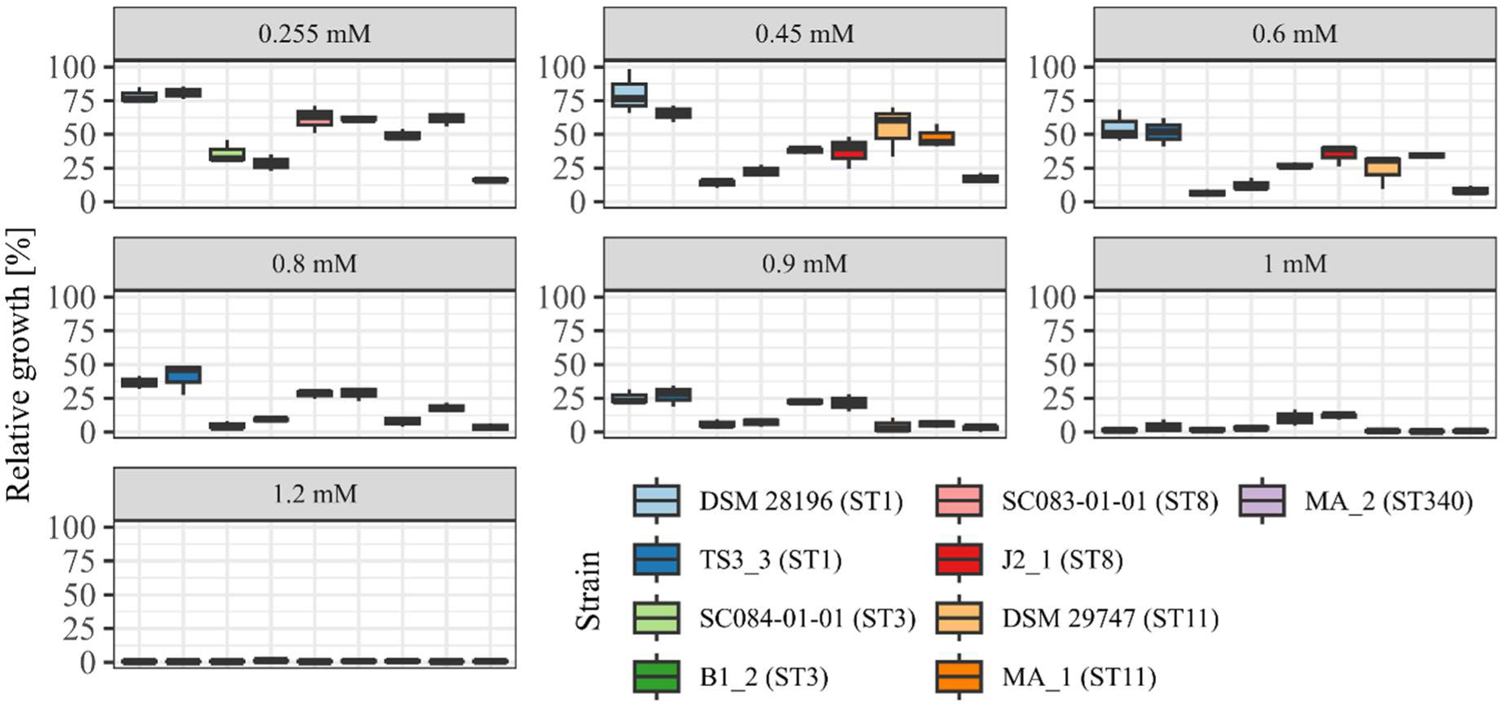
DCA tolerance at various concentrations. Tolerance of the strains to DCA was determined as relative growth compared to the reference culture (regarded as 100%) and MIC assay at concentrations from 0.255 mM to 1.2 mM. Strains of the same ST are depicted next to each other. No significant tolerance differences were observed within an ST at all concentrations.

### Sequencing-based assessments of prophage activity

Prophage prediction of all analyzed genomes exhibited various putative prophage regions, often with multiple incomplete and intact predicted regions in one genome (Table S1). Active prophages were determined by sequencing of particle-protected DNA. Sequencing reads were mapped to the corresponding host genome. Normalized read coverage (NPKM values) represented phage abundance as a relative measure indicative for phage activity (40). In the following, the term phage activity describes the production and resulting abundance of DNA-containing particles, and mentioned NPKM values refer to the OD-normalized data. Sequencing of phage DNA libraries was successful for all samples except for strain DSM 29747, which was the only strain without a predicted intact prophage genomic region (Table S1). This strain was therefore missing in further analyses.

Mapping of phage DNA sequencing reads onto respective host genomes is depicted in Figs 2 - 6. Overall examination of the mapping results revealed distinct activity of at least one region in all strains and under both induction conditions. This demonstrated spontaneous phage activity in all strains, and simultaneous activity of multiple phages within the same host. Almost all regions matched well with predicted and intact prophage regions. All these regions are summarized in Table 3 for name assignment to facilitate following descriptions. As apparent in Table 3, regions with positive prophage prediction accorded in size with typical genome sizes of *C. difficile* phages (41), while those without were significantly smaller.

**Fig 2.**
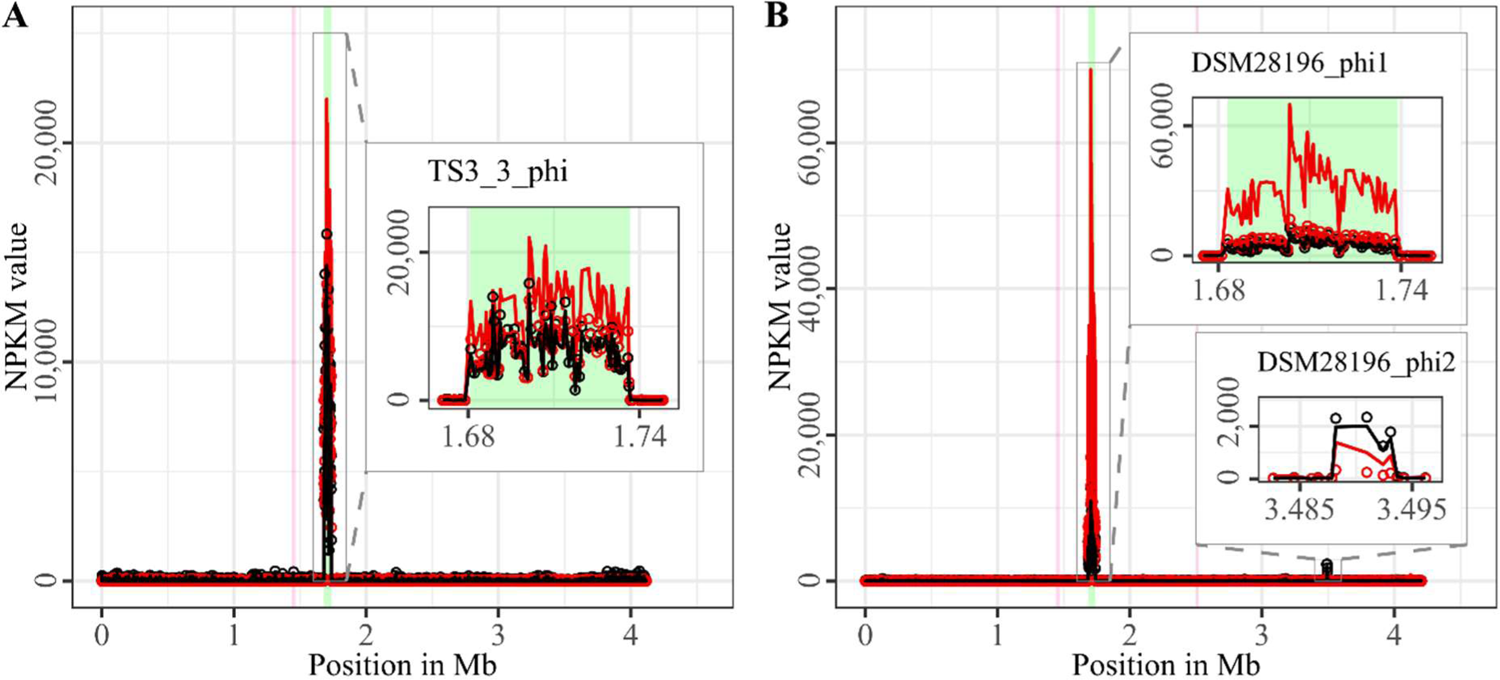
Coverage of phage sequencing reads of ST1-strains TS3_3 and DSM 28196. Sequencing reads of phage particles from spontaneous (black) and DCA-induced (red) release of strains (A) TS3_3 and (B) DSM 28196 were normalized to NPKM values (circles) with TraV (29), and NPKM values were additionally OD-normalized to OD_600_ = 2 (line graphs), before plotted against the chromosome (position in Mb) of the respective strain. Active regions were magnified for better visualization. Prophage regions predicted by PHASTEST (21) were highlighted in the background (intact = green, incomplete = pink).

**Table 3.**
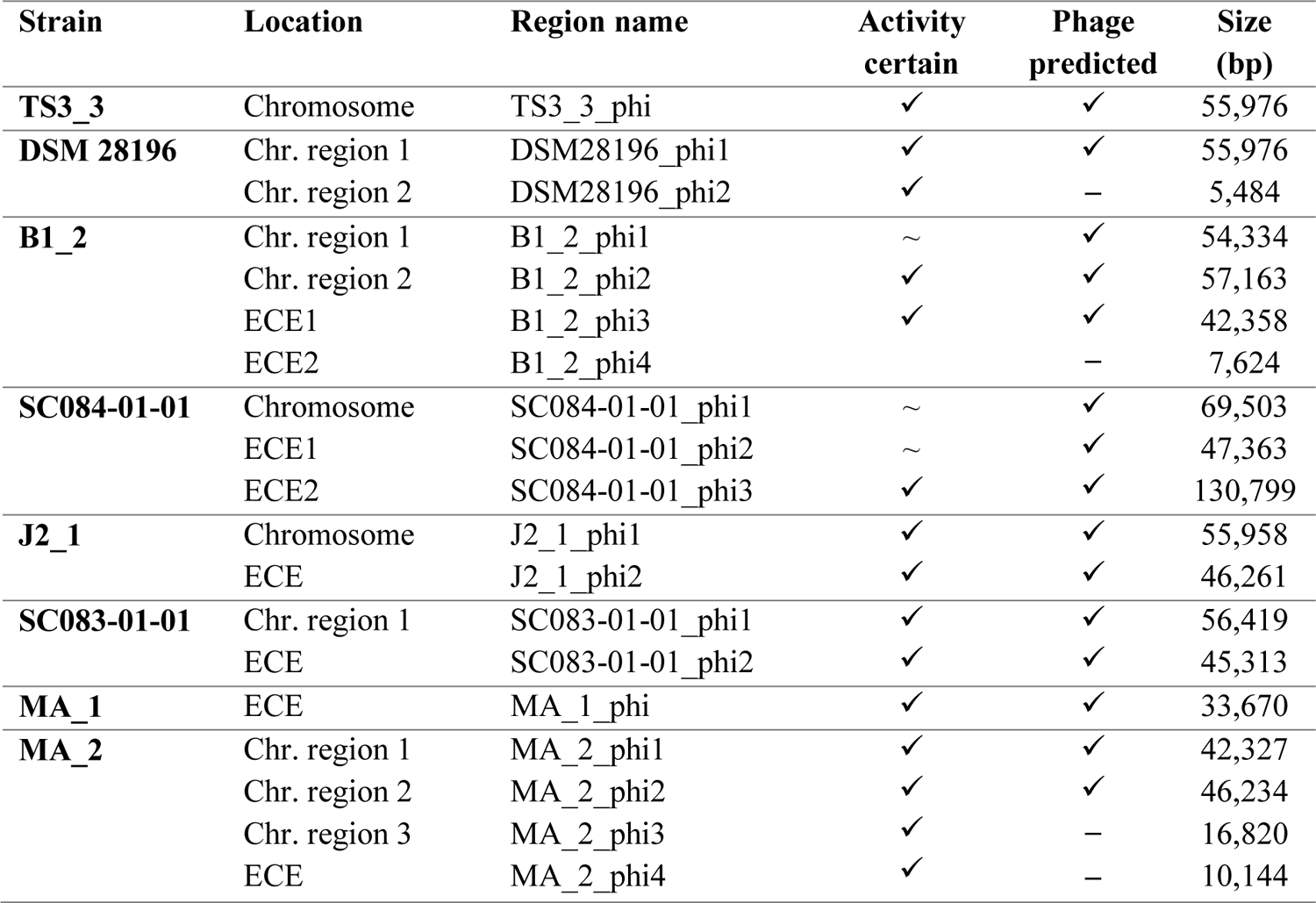
Overview of active regions in all strains. All active regions identified based on sequencing read coverage (Fig 2 - 6) were renamed according to a numbered scheme (strain_phiX). Regions were validated for certain (*✓*) or uncertain (∼) activity based on coverage signal strength. Additionally, prophage prediction with PHASTEST (21) is included by stating positive (*✓*) or negative (─) prediction, and region size in bp is stated as well.

Comparison of corresponding strains showed no apparent differences in carriage or location of active phages. Correspondingly, a correlation to the clinical background of a strain was not detected. As active prophages of corresponding strains resided at corresponding genome positions, we performed an ANIm analysis on extracted sequences of all active regions to assess their similarity amongst each other (Fig S1). This confirmed identical sequences of the analogous phages TS3_3_phi/DSM28196_phi1 of ST1-strains, and high similarity of both phages J2_1_phi1/SC083-01-01_phi1 and J2_1_phi2/SC083-01-01_phi2 among the ST8-strains, while the other phages exhibited only little or no similarity to the others.

All regions (Table 3) were further closely inspected their activities under DCA-induction. Overall NPKM-transformation to an OD of 2 revealed distinctly higher signals under DCA-induction in most active regions and all strains. This verified the phage-inducing effect of DCA, which varied apparently between the different regions even within the same host and thereby implied phage-dependency.

The genomes of both ST1-strains carried an identical phage (Fig S1) (TS3_3_phi/ DSM28196_phi1) as determined by ANIm analysis (Fig S1) and similar genome position. The analogous phages showed distinct spontaneous activity with approximate magnitude, but their DCA-induced activity differed substantially. Phage DSM28196_phi1 (Fig 2B) exhibited ∼3.5x higher signal increase under DCA than TS3_3_phi in the non-clinical strain (Fig 2A). Since the phages were identical, the differing reactions seemed to be host-related. The genome of strain DSM 28196 possessed another active region DSM28196_phi2, which showed minor activity under both conditions, and was phage-atypical by comprising only four genes and missing a prophage prediction (Fig 2 B).

The genomes of ST3-strains possessed several active regions (Fig 3), which were not similar to each other (Fig S1). In the genome of the non-clinical strain B1_2 (Fig 3A), two of the four active regions comprised the two ECEs. B1_2_phi4 on ECE2 exhibited the highest NPKM values under DCA-induction. Remarkably, B1_2_phi4 is another phage-atypical but active region without corresponding prophage prediction, as previously detected for DSM28196_phi2 (Fig 2B). B1_2_phi3 on ECE1 also showed increased activity under DCA, but both spontaneous and induced activity were not particularly high. B1_2_phi2 on the chromosome showed spontaneous activity and a substantial increase in DCA-induced activity. B1_2_phi1 on the chromosome exhibited almost no signal under spontaneous conditions, which slightly increased upon DCA-induction. This might indicate true DCA-induction of this phage without prior spontaneous activity. In contrast to strain B1_2, the corresponding clinical strain SC084-01-01 possessed no prominently active region on the chromosome (Fig 3B). Activity could be observed within the region SC084-01-01_phi1, but NPKM values were very low under both conditions and signals did not cover the whole phage region. Abundancy of this phage was probably insufficient to capture distinct activity by the sequencing approach. Similar activity was observed for SC084-01-01_phi2 on ECE1 of SC084-01-01. SC084-01-01_phi3 on ECE2 was the only region in SC084-01-01 with prominent activity under spontaneous conditions, and activity further increased apparently under DCA-induction.

**Fig 3.**
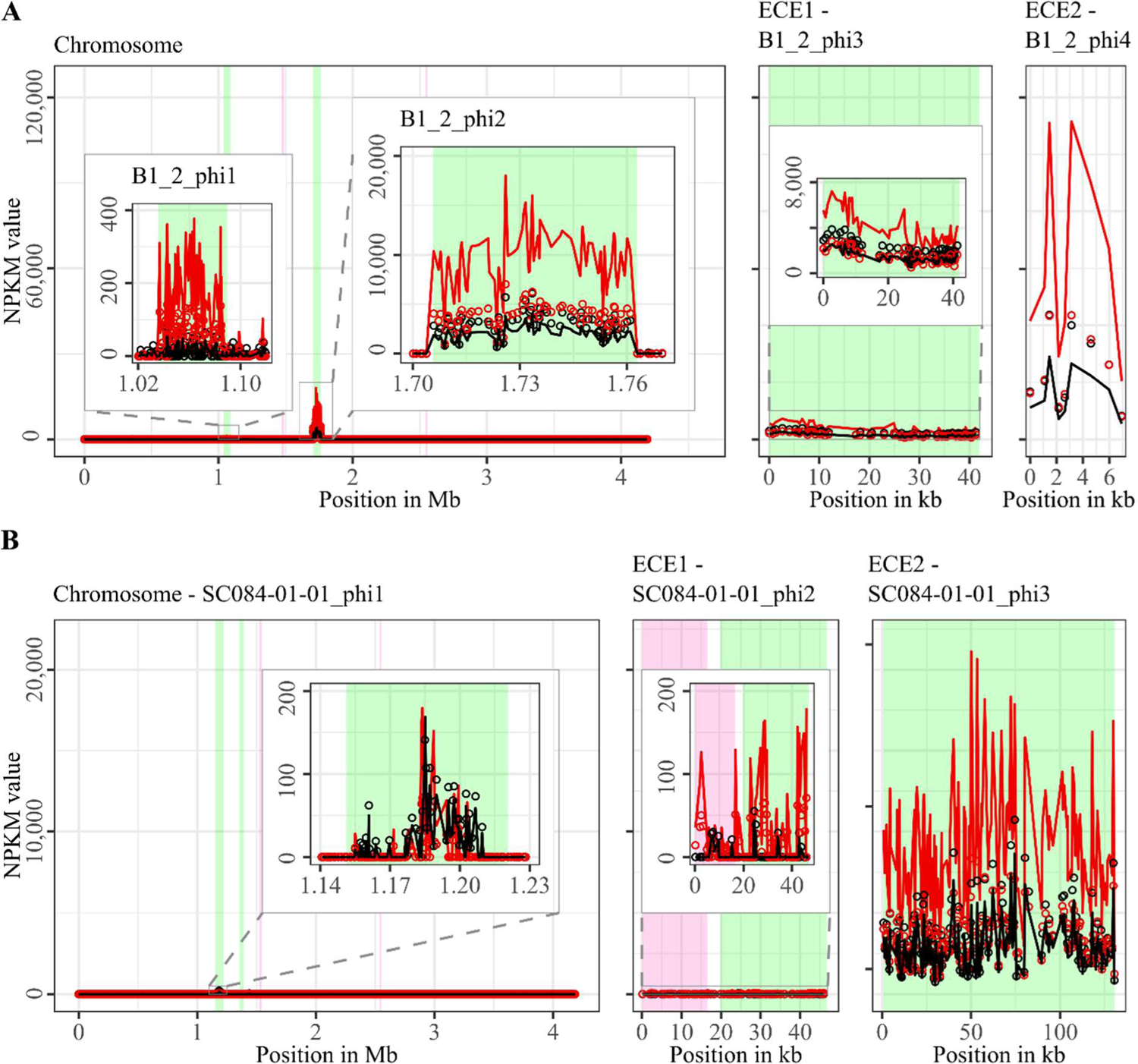
Coverage of phage sequencing reads of ST3-strains B1_2 and SC084-01-01. Sequencing reads of phage particles from spontaneous (black) and DCA-induced (red) release of strains (A) B1_2 and (B) SC084-01-01 were normalized to NPKM values (circles) with TraV (29), and NPKM values were additionally OD-normalized to OD_600_ = 2 (line graphs), before plotted against the chromosome (position in Mb) and ECE (position in kb) of the respective strain. Active regions were magnified for better visualization. Prophage regions predicted by PHASTEST (21) were highlighted (intact = green, incomplete = pink).

The genomes of both ST8-strains exhibited activity for their analogous chromosomal and extrachromosomal regions (Fig 4), which were similar phages according to ANIm analysis (Fig S1) and similar genomic location. J2_1_phi2 on the ECE of strain J2_1 showed distinct spontaneous activity and a DCA-induced activity increase (Fig 4A), whereas SC083-01-01_phi1 on the ECE of SC083-01-01 was only slightly active under spontaneous conditions, and signal increase upon DCA-induction was only little. The activity of the chromosomal phages differed apparently as well. J2_1_phi1 on the J2_1 chromosome was spontaneously active and showed increased activity under DCA-induction (Fig 4A). The left part of the prophage region started with minor activity, which drastically increased at the terminase genes. Strikingly, the left part of the prophage region started with minor activity, which drastically increased at the terminase genes. Further remarkable, sequencing reads mapped beyond the predicted prophage region and spread upstream (∼30 kb) and downstream (∼135 kb). The downstream region adjoined the phage activity with similar NPKM values that gradually decreased over the entire section. The chromosomal phage SC083-01-01_phi1 of the corresponding clinical ST8-strain SC083-01-01 did not exhibit these peculiarities (Fig 4B). It was spontaneously active, and activity increased substantially under DCA treatment. This increase was strikingly twice as high as observed for the analogous phage J2_1_phi1 (Fig 4A), despite their similarity (Fig S1). Such dissimilar activity increase among analogous phages was already observed in the ST1 strains (Fig 2). In both ST8- and ST1-pairs, a distinctly stronger increase upon DCA-induction was connected to the clinical background of the strains. Since DCA-stress levels did not significantly differ between corresponding clinical and non-clinical strains (Fig 1) and thereby suggested similar induction levels, the question for the underlying mechanism of these diverging activities in analogous phages arose. This prompted to an undefined regulation of phage induction involved in the clinical strains.

**Fig 4.**
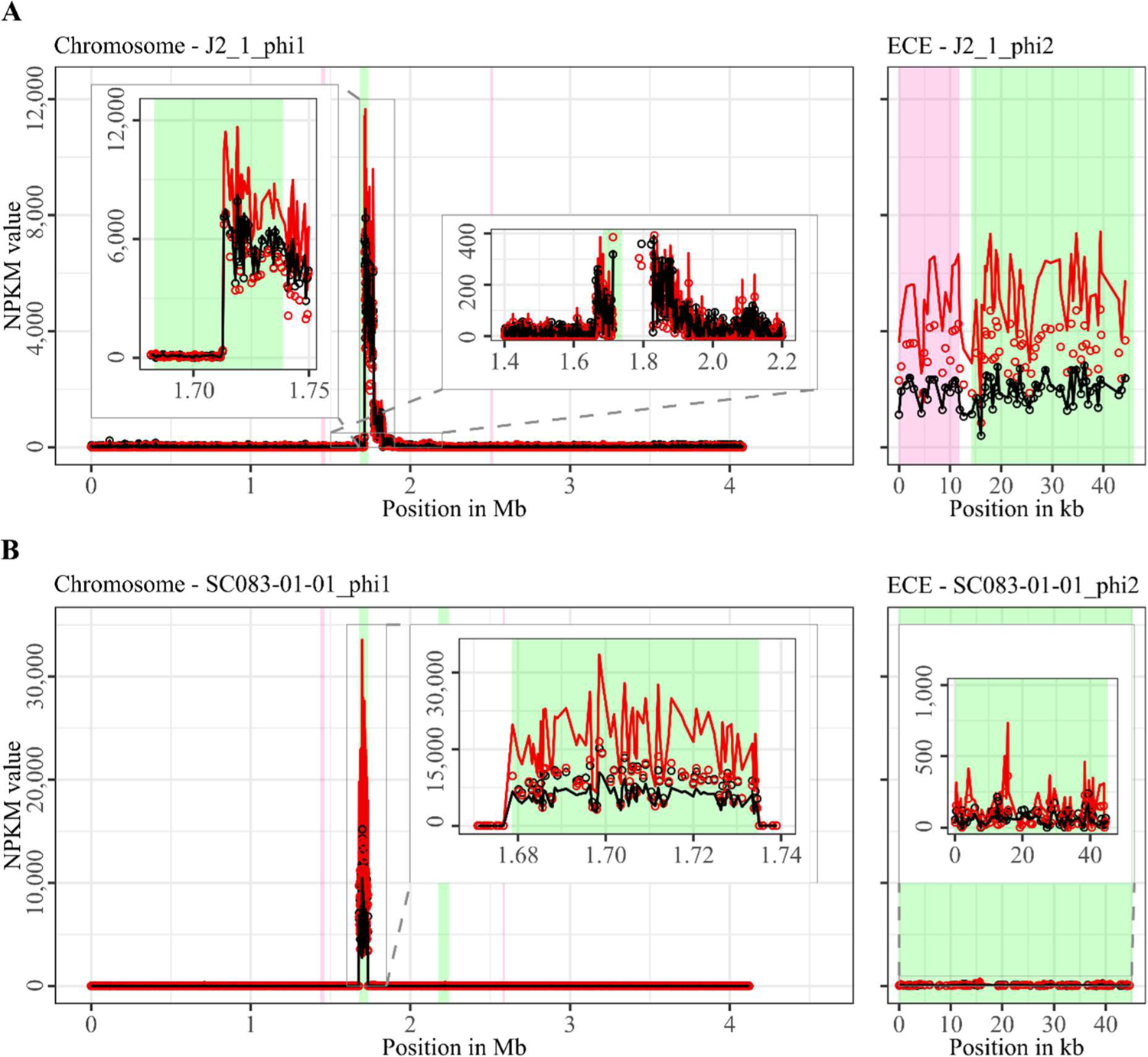
Coverage of phage sequencing reads of ST8-strains J2_1 and SC083-01-01. Sequencing reads of phage particles from spontaneous (black) and DCA-induced (red) release of strains (A) J2_1 and (B) SC083-01-01 were normalized to NPKM values (circles) with TraV (29), and NPKM values were additionally OD-normalized to OD_600_ = 2 (line graphs), before plotted against the chromosome (position in Mb) and ECE (position in kb) of the respective strain. Active regions were magnified for better visualization. Prophage regions predicted by PHASTEST (21) were highlighted (intact = green, incomplete = pink).

Strain MA_1 could not be compared to its corresponding clinical strain DSM 29747, but distinct spontaneous activity was visible for MA_1_phi on the ECE, which increased under DCA treatment (Fig 5).

**Fig 5.**
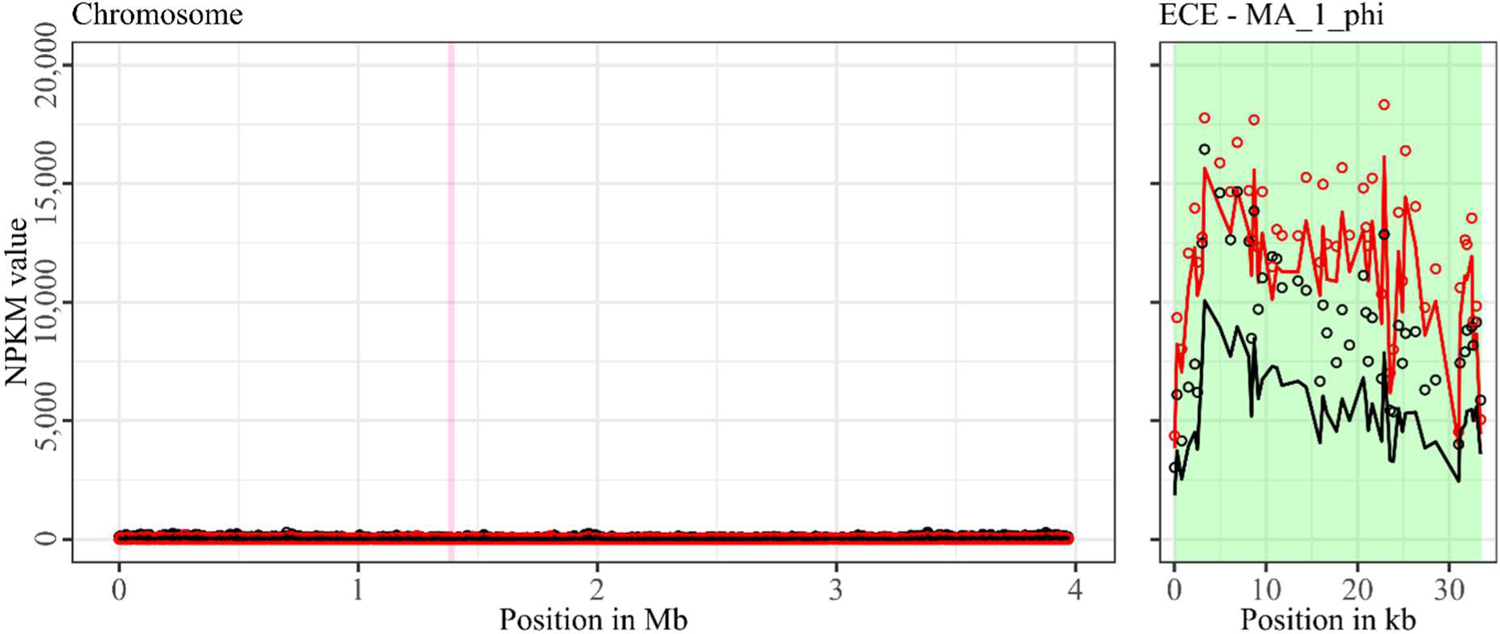
Coverage of phage sequencing reads of ST11-strain MA_1. Sequencing reads of phage particles from spontaneous (black) and DCA-induced (red) release of strain MA_1 were normalized to NPKM values (circles) with TraV (29), and NPKM values were additionally OD-normalized to OD_600_ = 2 (line graphs), before plotted against the chromosome (position in Mb) of MA_1. Active regions were magnified for better visualization. Prophage regions predicted by PHASTEST (21) were highlighted (intact = green, incomplete = pink).

Strain MA_2 is to our knowledge the first cryptic *C. difficile* strain with detailed phage examination. This strain possessed four active regions (Fig 6). MA_2_phi2 on the chromosome and MA_2_phi4 on the ECE both showed prominent activity under spontaneous conditions and a distinct activity increase upon DCA-induction. Interestingly, MA_2_phi4 was another active region without phage prediction, as observed previously for DSM28196_phi2 (Fig 2B) and B1_2_phi4 (Fig 3A). This was also true for MA_2_phi3 on the chromosomal, where activity was however very low under spontaneous conditions. Activity of this region increased upon DCA-induction. Chromosomal region MA_2_phi1 exhibited least activity in this genome even under DCA treatment. Read mapping for MA_2_phi1 and MA_2_phi2 did not cover the entire predicted prophage regions, but sections without read coverage contained only bacterial genes and were therefore evidently mis-predicted.

**Fig 6.**
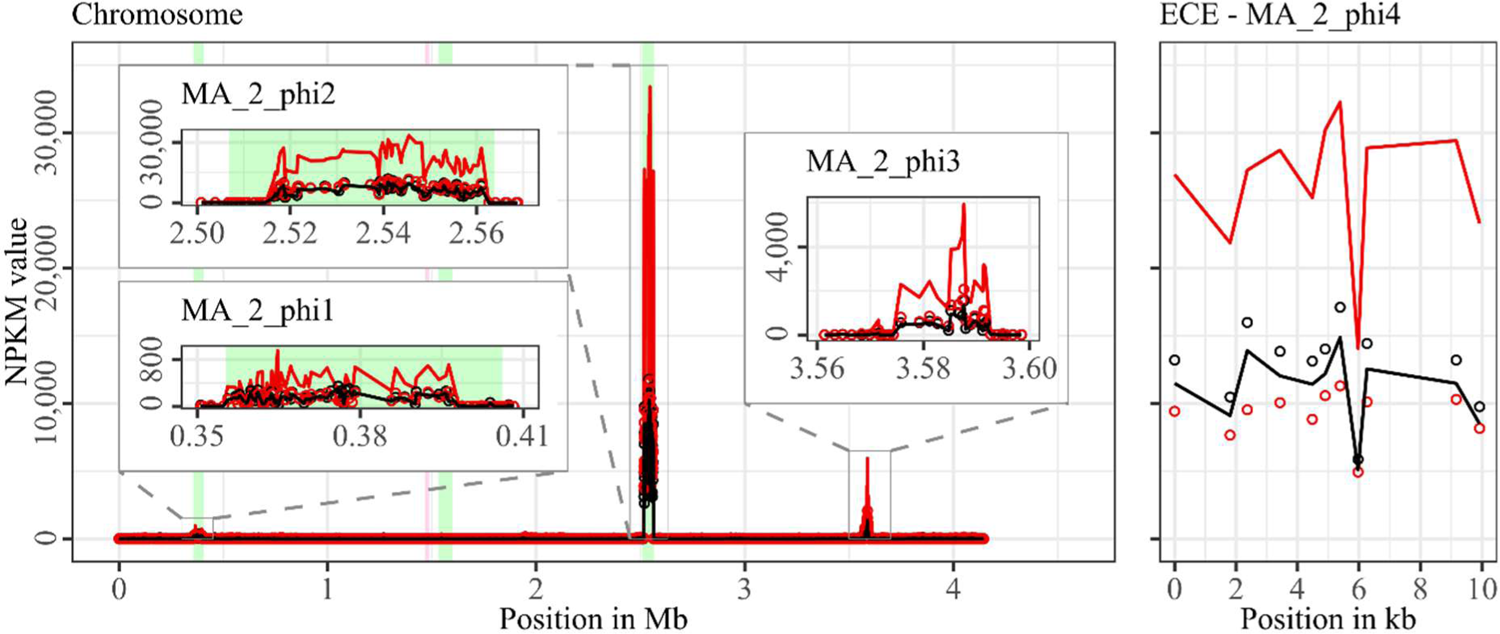
Coverage of phage sequencing reads of cryptic ST340-strain MA_2. Sequencing reads of phage particles from spontaneous (black) and DCA-induced (red) release of strain MA_2 were normalized to NPKM values (circles) with TraV (29), and NPKM values were additionally OD-normalized to OD_600_ = 2 (line graphs), before plotted against the chromosome (position in Mb) and ECE (position in kb) of MA_2. Active regions were magnified for better visualization. Prophage regions predicted by PHASTEST (21) were highlighted (intact = green, incomplete = pink).

Almost all identified active prophage regions could be induced by the secondary bile salt DCA. Mentionable, assessed phage activity after DCA treatment might be influenced by a direct effect of DCA on the phage. A study on different bacteriophages in *Escherichia coli* investigated the effect of bile salts on the host-phage interaction and observed varying survival rates of the phages (42).

Moreover, almost all ECEs were detected as active phages. Although extrachromosomal prophages have already been described in *C. difficile* strains (43,44), only a few of these were isolated and characterized so far (41).

### Phage genome annotation and gene content analysis

#### Phage genomes harbor virulence-relevant genes

All active regions identified via sequence read mapping (Figs 2 – 6, Table 3) were inspected after anew annotation with Pharokka (32) for genes that might increase virulence of the host (genomic information in supplementary data file S1). All of them exhibited characteristic phage genes in a modular organization according to the different encoded functions, as typically seen in *C. difficile* phages (16,43,45). In some genomes, proteins characteristic for plasmids were found, such as genes encoding partition proteins and replication initiation factors (46). Genes encoding plasmid-related proteins besides phage-typical ones are common in a certain type of MGE, so-called phage-plasmids (47). These phage-plasmid features were especially recorded for extrachromosomal prophages, but were also present on the chromosomally integrated prophage MA_2_phi1. Further frequently observed genes in the phage genomes encoded proteins with potential involvement in cellular metabolism and growth, such as metallo-proteases, kinases, a phosphatase, and most of all putative rhodanese-related sulfurtransferases. These genes might be advantageous for the bacterial fitness, thereby indirectly contributing to host virulence. Two phage genomes (B1_2_phi1 and SC084-01-01_phi1) carried genes encoding hemolysin XhlA, an established virulence factor in other bacterial species (48), capable of lysing mammalian erythrocytes (49). In the opportunistic pathogen *Mannheimia haemolytica*, temperate phages were induced that encoded hemolysin XhlA and discussed to contribute to bacterial pathogenicity and transfer of this virulence factor (50). Hemolysis in *C. difficile* is not commonly known, but few studies demonstrated its hemolytic capability (51). However, XhlA is also present in other prophage genomes as part of the lysis module (52). Indeed, the gene encoding XhlA was found next to the lysis-relevant genes encoding holin and endolysin in phages B1_2_phi1 and SC084-01-01_phi1. Thus, the actual role of hemolysin XhlA in these phages and its potential impact on host virulence remains unclear. Phage SC084-01-01_phi1 possessed a gene encoding an ABC-transporter, which might contribute to antibiotic resistance of the host (53). Phage SC084-01-01_phi3 harbored a putative spore protease, which could influence the bacterial sporulation or germination ability and, as a consequence, alter bacterial fitness (43). Other genes conferring antibiotic resistances or encoding known virulence factors were not identified in the active phage genomes. Six phages (TS3_3_phi, DSM28196_phi1, B1_2_phi2, J2_1_phi1, SC083-01-01_phi, MA_2_phi2) carried arrays of clustered regularly interspaced short palindromic repeats (CRISPR), which is similar to other *C. difficile* phages with described CRISPRs (37,43,54). Temperate phages carrying CRISPR arrays increase host immunity against other invading phages (55). The corresponding host genomes were verified to encode Cas proteins required for CRISPR-Cas-mediated phage immunity (56). CRISPRs in prophages represent horizontally transferrable immunity against phages, which is specifically relevant in the context of phage therapy, an alternative treatment approach for bacterial infections with growing importance in view of increasing multidrug-resistances (57).

#### Active non-phage elements likely belong to so far undescribed HGT mechanisms in *C. difficile*

The regions without corresponding prophage prediction (Table 3) did not possess a phage-typical genome accordingly. Therefore, further gene analysis of these non-phage elements were performed based on the original genome annotation with Prokka (58) (genomic information in supplementary data file S1). No proteins involved in capsid or tail production, DNA packaging or host lysis were present. The lack of structural proteins was striking, since these DNA elements were enveloped according to the DNA isolation procedure. This indicated the involvement of unrelated particles. Even additional analysis of hypothetical proteins with InterProScan (34) and BLASTp (33) could not identify further functions. All other proteins were assigned to functions with DNA-binding activity, like helicases, integrases, relaxases, and transcriptional regulators. These genes are typical for phage genomes but also for MGEs like transposons as Integrative and Conjugative or Mobilizable Elements (ICE / IME) (59). Screening for these MGEs with ICEscreen (60) validated DSM28196_phi2 as complete IME, while MA_2_phi3 was detected as invalid ICE. Interestingly, the peculiar upstream-region of J2_1_phi1 (Fig 4A) was also identified as an ICE, although being incomplete. These integrative MGEs do not encode proteins for production of particles that carry the respective mobile sequence (59). Interestingly, transposons were found to hitchhike co-residing phages in several bacteria such as *Staphylococcus aureus* (61), *Vibrio cholerae* (62), and *Enterococcus faecalis* (63), enabling the phage-mediated transduction of virulence-relevant genes. This type of transduction was demonstrated once in *C. difficile* with the transfer of a conjugative transposon carrying an antibiotic resistance gene (10). Transduction can be either generalized, specialized or lateral (64). They all imply the “headful” DNA-packaging, in which the terminase starts DNA packaging at a bacterial homologue to the phage packaging site until the capsid is full, which consequently implies that transduced DNA is at least of similar phage genome size (64). The transduction mechanisms differ in transduced DNA and frequency (65). Specialized and lateral transduction involve host DNA adjacent to the prophage, while random host DNA is packaged in generalized transduction (65). Generalized and specialized transduction are processes of erroneous DNA-packaging, which results in low transduction frequency detectable by sequencing read coverage (64,65). In contrast, lateral transduction results in high sequencing read coverage comparable to actual phage activity, as this mechanism is assumed as natural phage trait instead of mistaken processes (64,65). This trait comprises phage genome replication and simultaneous DNA-packaging before excision from the chromosome, whereby a substantial amount of adjacent host DNA is packaged as well (66,67). All these characteristics of lateral transduction accorded with the observed mapped region downstream of phage J2_1_phi1 (Fig 4A), which indicated involvement of this DNA segment in phage-mediated lateral transduction.

The downstream region of J2_1_phi1 (Fig 4A) did not comprise characteristic genes for MGEs. Instead, several genes encoded proteins with potential relevance for strain virulence, such as genes for ABC-transporters, a multidrug efflux system ATP-binding protein, stress-related proteins, proteins involved in resistance to vancomycin and daunorubicin, and the putative virulence factor BrkB. Therefore, mobilization and transfer of this region is critical regarding the spread of antibiotic resistances or virulence-related genes. The drastic NPKM difference within the genome phage J2_1_phi1 (Fig 4A) might also result from the process of lateral transduction, as inaccurate excision of the phage genome after *in situ* replication leads to phage genome truncation.

Since no evidence for lateral transduction was observed for phage J2_1_phi1’s analogue SC083-01-01_phi1 (Fig 4B) despite their high similarity (Fig S1), the question about underlying differences arose. Direct phage genome comparison revealed diverging excisionases and integrases as well as four additional amino acids in the large terminase protein of J2_1_phi1. All these proteins perform activities destining for lateral transduction (64).

The extrachromosomal non-phage elements were significantly smaller than the co-existing phages (Table 3), which is in contrast with the headful packaging mechanism required in transduction. This indicate a form of DNA-protecting particle other than phages, such as gene transfer agents (GTA). These phage-like particles carry DNA between 4 to 14 kb (68), which is similar to the sizes of the detected non-phage elements (Table 3). However, GTAs package bacterial DNA randomly (68), which does not fit to the detected distinct activity of specific regions, making GTAs unlikely as mode of action. Extracellular vesicles are known in various bacteria and described to carry and transfer genetic content, e.g. plasmids, between cells (69–71). This type of alternative HGT is not well characterized so far, but was demonstrated to allow interspecies gene transfer (71), which underlines the importance of vesicle-mediated DNA exchange. Noteworthy, vesicle-driven HGT in *C. difficile* has not been described so far.

### Classification of the active phages

#### Terminase-based determination of the phage DNA-packaging strategy

Assessment of the phage-DNA packaging mechanisms was performed to validate the above hypothesized transduction events. The large terminase subunit was analyzed via protein sequence alignment and phylogenetic tree construction referring to Rashid et al. (37). This assigned the phages to different phage DNA-packaging mechanisms (Fig 7). All our phages were assigned to clusters comprising other *C. difficile* phages, which predominantly represented the P22-like headful packaging mechanism, followed by the 3’-extended COS ends and an unknown strategy. Consequently, most of the phages were predicted to utilize the headful packaging mechanism and would, thus, be indeed capable of transducing host DNA. The mechanism “P22-like headful” originates from the packaging strategy employed by phage P22 of *Salmonella enterica*. Phage P22 was originally described to perform generalized transduction (72), but recent evidence demonstrated also specialized as well as lateral transduction activity (67). These terminase analysis results supported the assumption of lateral transduction of the phage J2_1_phi1 downstream region, which is by that the first described case in *C. difficile*.

**Fig 7.**
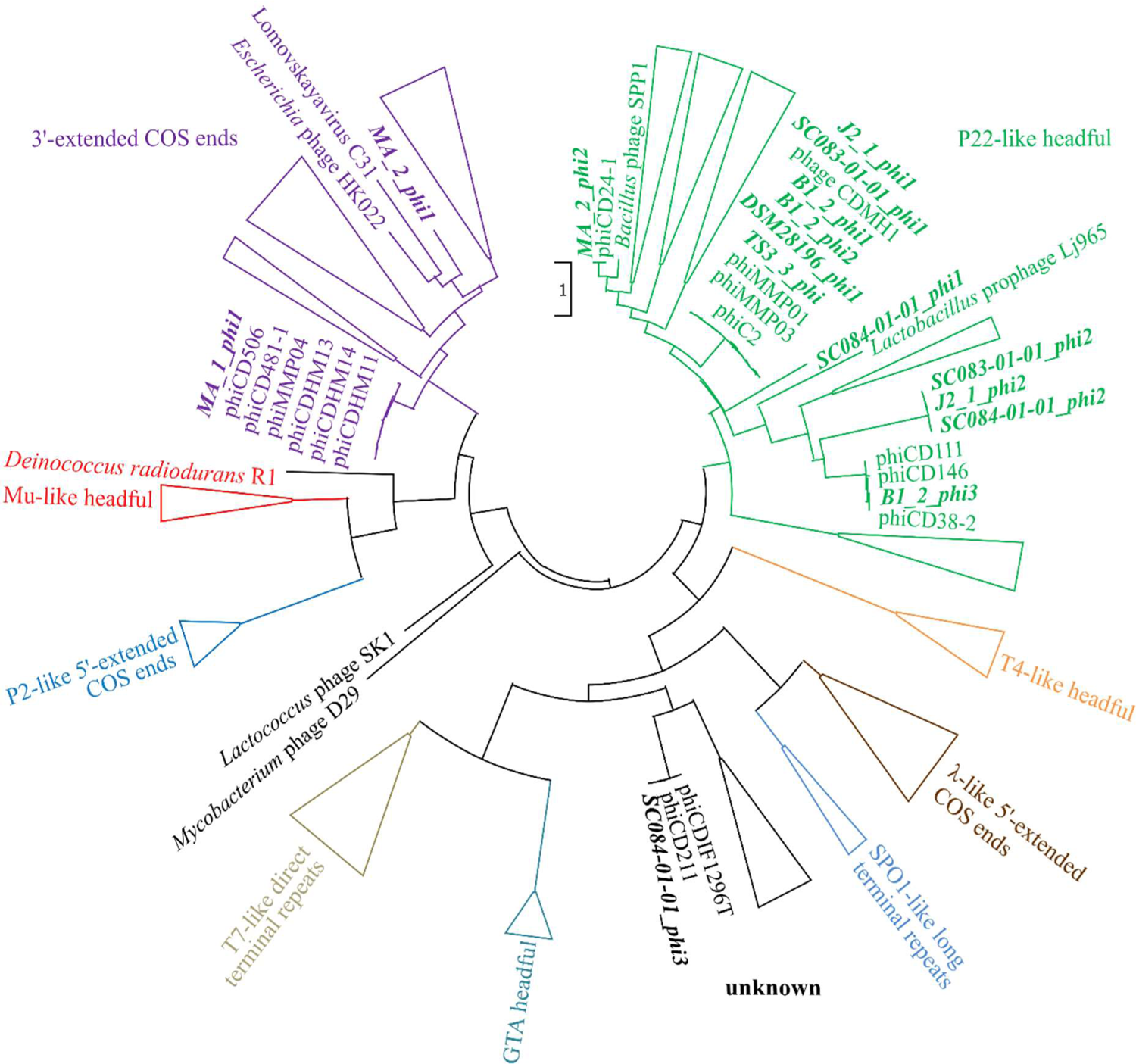
Maximum-likelihood phylogenetic tree of large terminase protein sequences. The large terminase of the active phages (highlighted in bold and italic) were aligned on protein level to the reference sequences from Rashid et al. (37). Branches of the different DNA-packaging strategies were colored according to Rashid et al. (37), and branches were collapsed for better visualization if none of our phages was included.

#### Nucleotide BLAST analyses assess phage prevalence and novelty

A nucleotide BLAST analysis (33) was performed on all active regions in Table 3 to check for similar phages and elements, and to assess prevalence among *C. difficile* strains. The results are available in supplementary data file S2 and summarized in Table 4. Most of the phages matched against *C. difficile* phages with query coverages between 4% and 90% and percent identities between 86.26% and 99.86%. This confirmed that our phages are indeed similar to known phages but still represent novel types, which underlines the contribution of this work to the general knowledge on *C. difficile* phages. Further, chromosomal phages also matched against a multitude of *C. difficile* chromosomes, while the extrachromosomal phages often corresponded to *C. difficile* plasmids and few chromosomes. This demonstrated the prevalence of the identified phages in other *C. difficile* strains. The non-phage elements B1_2_phi4 and MA_2_phi4 yielded no significant BLAST hit against a phage, but matched against *C. difficile* plasmids. Both matching plasmids belong to classes of *C. difficile* plasmids with similar organization that are present in diverse *C. difficile* strains (73,74). All these plasmids might therefore likewise be inducible and particle-protected, which implies a different mechanism of HGT than currently assumed.

**Table 4.**
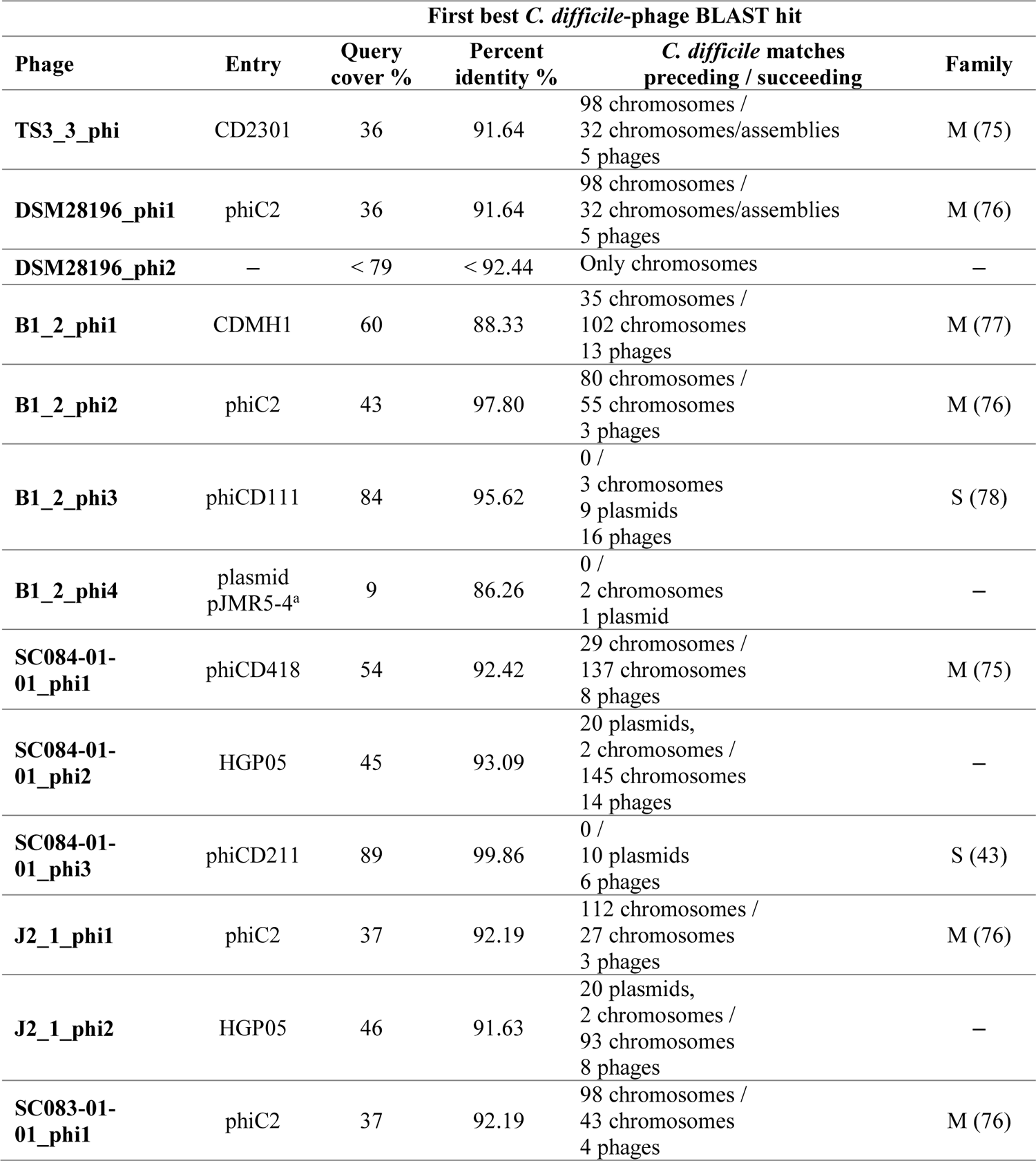
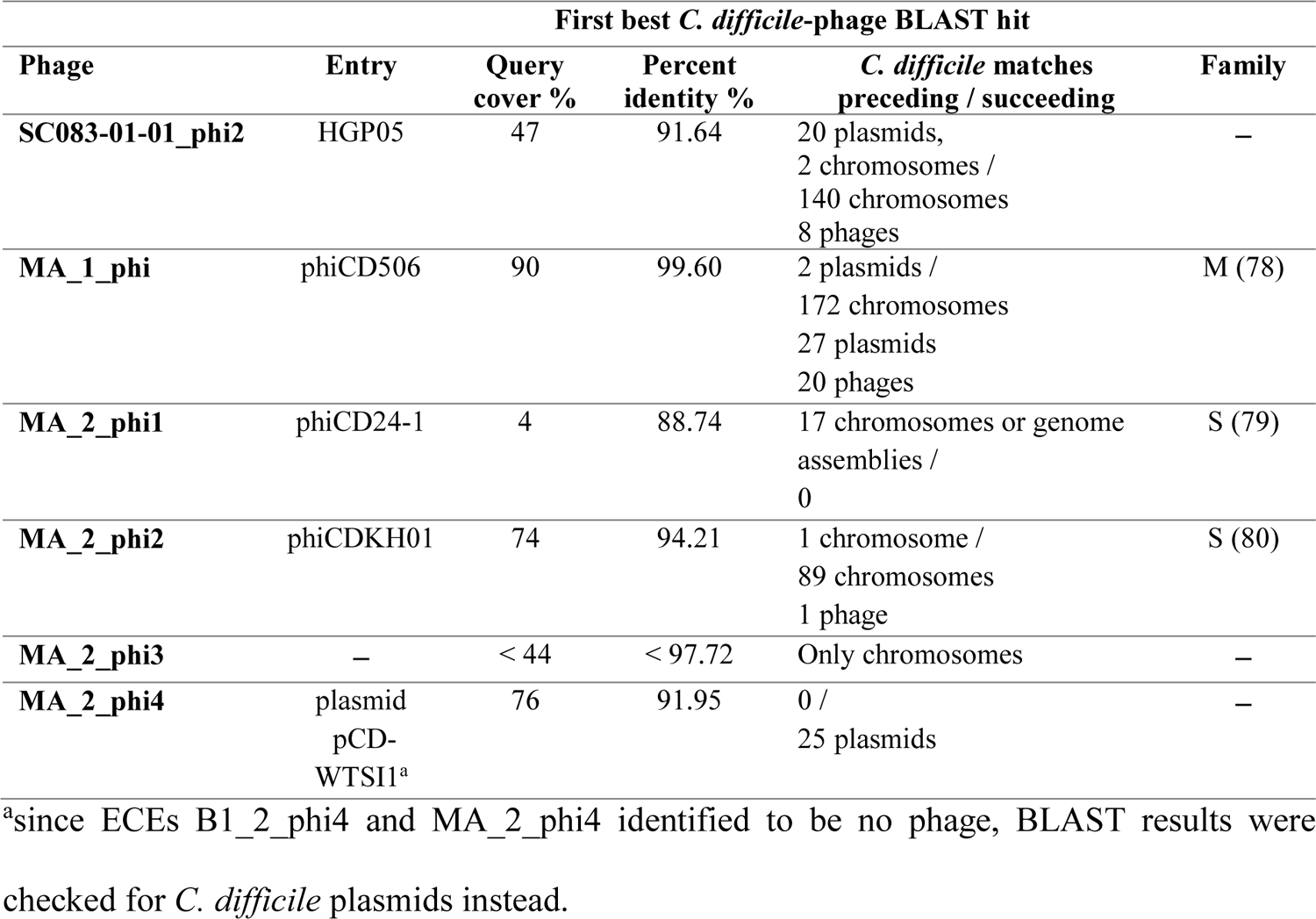
Nucleotide BLAST results of the identified active regions. Results were summarized regarding the first phage-related *C. difficile* BLAST hit by stating its description, query coverage, percent identity, and number of pre- and succeeding chromosome/plasmid entries. The few hits of metagenome-assembled phages were not listed. If available, information on the phage family of the respective hit is indicated (M – *Myoviridae*, S – *Siphoviridae*).

### Genome-based phage assignment to *Myoviridae* and *Siphoviridae*

We lastly classified our phages morphologically. All known *C. difficile* phages so far belong to the *Caudovirales* family of *Myoviridae* or *Siphoviridae*, which distinguish by tail appearance (41). Genome inspection for the presence of baseplate proteins characteristic for *Myoviridae* and the length of the tail length tape measure protein as indicator for *Siphoviridae* could assign eleven phages to *Myoviridae* and four phages to *Siphoviridae* (Table S2).

## Conclusion

We aimed to investigate prophage activity in different clinical and non-clinical *C. difficile* strains and unravelling potential relationships between phage activity and clinical background of the strain. Our investigations did not find specific connections to the clinical background, although we observed stronger DCA-related activity with clinical background for phages that were similar between clinical and non-clinical strains. We further revealed several interesting findings with relevance for future *C. difficile* phage research. We identified and characterized several active prophages in various *C. difficile* strains with a sequencing-based approach. This sensitive approach allowed detecting multiple co-existing prophages with diverse activity. Most of these phages were distinctly active without specific induction, but they showed increased activity when induced with the secondary bile salt DCA. This proved that spontaneous activity is common in *C. difficile* prophages, and that the natural stressor DCA triggers prophage induction. These findings are crucial for investigating *C. difficile* biology, since phages evidently affect *C. difficile* fitness and virulence by influencing toxin production or participating in the exchange of clinically relevant genes. We also found such genes with potential connection to virulence in some phage genomes. In this context, research on actual *in vivo* phage mobility should increasingly resemble *C. difficile*’s natural habitat. The sequencing approach further revealed active regions without phage identity. Based on genomic examinations, these regions were identified as another form of MGE, in most cases possibly integrative elements. These elements apparently participated in a strategy of mobilization that involves some kind of DNA envelopment, which pointed to phage particles or bacterial vesicles. This phenomenon was observed in several of the analyzed strains, which indicated that this type of DNA mobilization might be a frequent mechanism in *C. difficile* and should be further investigated. We also observed transduced bacterial DNA that was likely the result of lateral transduction employed by one phage, which has not been described so far in *C. difficile* and opens up new perspective on *C. difficile* phage research.

## Acknowledgements

We thank Melanie Heinemann for technical assistance. This work was funded by the Federal State of Lower Saxony, Niedersächsisches Vorab CDiff and CDInfect projects (VWZN2889/3215/3266). We acknowledge support by the Open Access Publication Funds of the Göttingen University. This work was partly supported by the Göttingen Graduate Center for Neurosciences, Biophysics, and Molecular Biosciences at the Georg-August-Universität Göttingen. The funders had no role in study design, data collection and analysis, decision to publish, or preparation of the manuscript.

## Notes

### Competing Interest Statement

The authors have declared no competing interest.

### Summary of Updates

We added a sentence in the Acknowledgments section and changed the IDs of 2 strains in a supplementary file

