## Supplementary figure S1 for "Novel insights into phage biology of the pathogen *Clostridioides difficile* based on the active virome"

### Supplementary material Fig S1

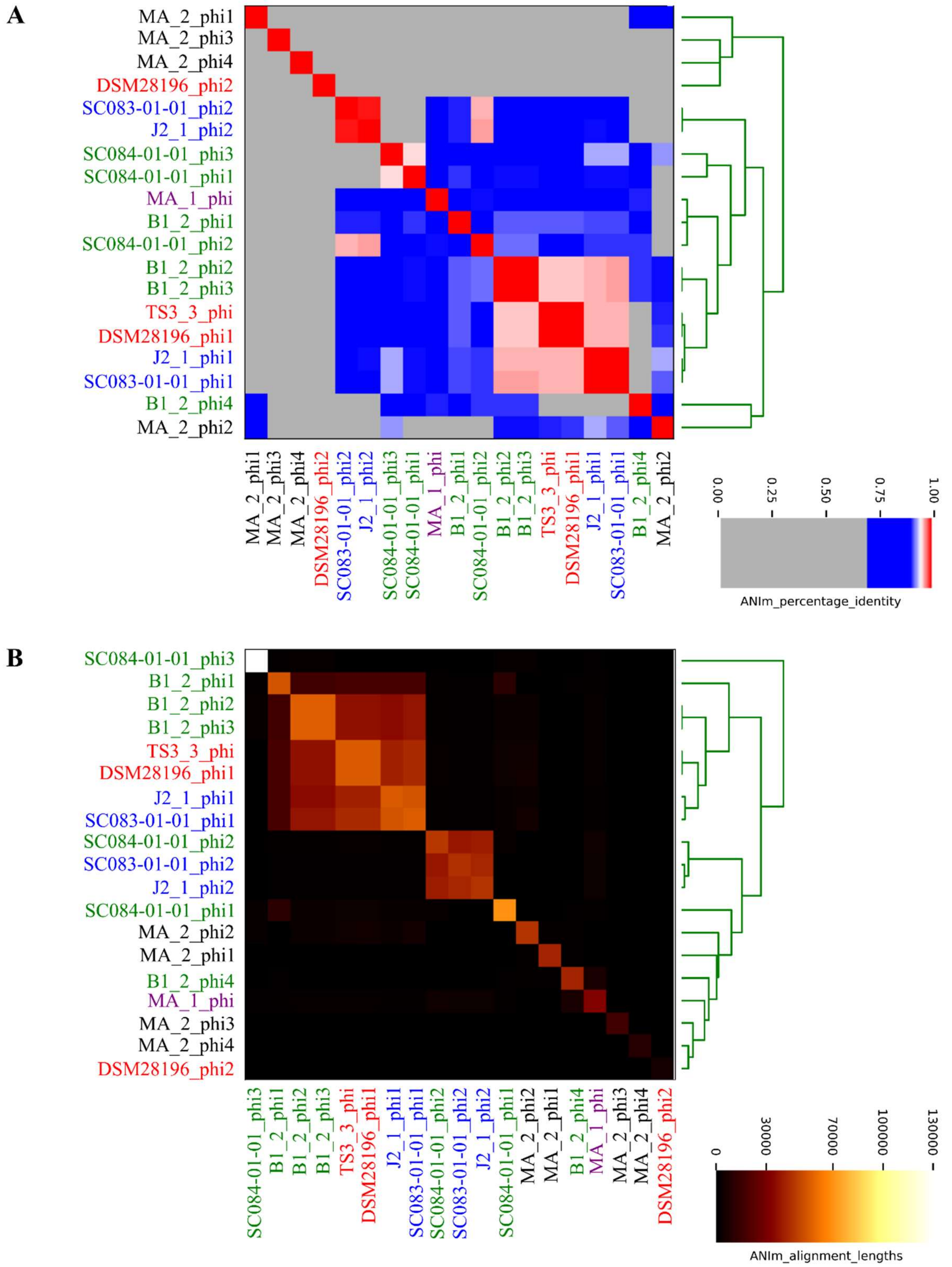


**Fig S1. ANIm analysis of the active prophage regions.** Heatmaps depict the (A) ANI values and (B) alignment lengths among the various active regions. The active regions are color-coded according to their ST for comparison better of analogous phages: red = ST1, green = ST3, blue = ST8, purple = ST11, black = ST340.
