## Supplementary table S1 and S2 for "Novel insights into phage biology of the pathogen *Clostridioides difficile* based on the active virome"

**Supplementary material Table S1 & S2**

**Table S1. Detailed prophage prediction results.** More detailed PHASTEST (1) results of the analyzed strains on predicted regions and completeness score.

| Strain | Predicted region | completeness |
| --- | --- | --- |
| *C. difficile* TS3_3 | 1,437,930 – 1,465,233  1,680,766 – 1,736,741 | incomplete  intact |
| *C. difficile* B1_2  Chromosome  ECE 1  ECE 2 | 1,035,372 – 1,089,705  1,467,664 – 1,488,405  1,705,680 – 1,762,842  2,540,996 – 2,554,708  19 – 41,921  none | intact  incomplete  intact  incomplete  intact |
| *C. difficile* J2_1  Chromosome  ECE | 1,439,999 – 1,467,302  1,682,838 – 1,738,813  2,503,885 – 2,518,076  1 – 11,796  14,174 – 46,002 | incomplete  intact  incomplete  incomplete  intact |
| *C. difficile* MA_1  Chromosome  ECE | 1,375,651 – 1,402,933  28 – 33,560 | incomplete  intact |
| *C. difficile* MA_2  Chromosome  ECE | 355,247 – 406,180  1,466,266 – 1,487,446  1,532,128 – 1,599,211  2,506,900 – 2,563,763  none | intact  incomplete  intact  intact |
| *C. difficile* DSM 28196 | 1,439,999 – 1,467,302  1,682,838 – 1,738,813  2,503,885 – 2,518,076 | incomplete  intact  incomplete |

**Table S1 continued.**

| Strain | Predicted region | completeness |
| --- | --- | --- |
| *C. difficile* SC084-01-01  Chromosome  ECE 1  ECE 2 | 1,151,245 – 1,220,747  1,357,429 – 1,392,299  1,521,667 – 1,542,408  2,537,975 – 2,551,687  136 – 16,524  19,825 – 46,846  736 – 130,763 | intact  intact  incomplete  incomplete  incomplete  intact  intact |
| *C. difficile* SC083-01-01  Chromosome  ECE | 1,434,660 – 1,462,107  1,678,611 – 1,735,029  2,173,498 – 2,243,863  2,578,831 – 2,592,871  58 – 45,180 | incomplete  intact  intact  incomplete  intact |
| *C. difficile* DSM 29747 | 1,375,725 – 1,403,007 | incomplete |

**Table S2. Genome-based prediction of morphological family.** Sequence length of the tail tape measure protein and presence of baseplate proteins were used to predict a phage of belonging to the *Myo-* (M) or *Siphoviridae* (S).

| Phage | Tail length tape measure protein length (aa) | Presence Baseplate | Family prediction |
| --- | --- | --- | --- |
| TS3_3_phi | 770 | + | M |
| DSM28196_phi1  DSM28196_phi2 | 770  ─ | +  ─ | M  ─ |
| B1_2_phi1  B1_2_phi2  B1_2_phi3  B1_2_phi4 | 764  797  1,767  ─ | +  +  ─  ─ | M  M  S  ─ |
| SC084-01-01_phi1  SC084-01-01_phi2 SC084-01-01_phi3 | 1,416  1,130  2,000* | +  +  ─ | M  M  S |
| J2_1_phi1  J2_1_phi2 | 797  1,129 | +  + | M  M |
| SC083-01-01_phi1  SC083-01-01_phi2 | 797  1,129 | +  + | M  M |
| MA_1_phi | 584 | + | M |
| MA_2_phi1  MA_2_phi2  MA_2_phi3  MA_2_phi4 | 2,226  1,839  ─  ─ | ─  ─  ─  ─ | S  S  ─  ─ |

*SC084-01-01_phi1 possessed two putative tail length tape measure proteins in close proximity. This is similar to the recent entry of phiCD211 (NC_029048.2), whereas the smaller protein is annotated as minor tail protein in the old entry LN681537.2.
